# IDseq – An Open Source Cloud-based Pipeline and Analysis Service for Metagenomic Pathogen Detection and Monitoring

**DOI:** 10.1101/2020.04.07.030551

**Authors:** Katrina L. Kalantar, Tiago Carvalho, Charles F.A. de Bourcy, Boris Dimitrov, Greg Dingle, Rebecca Egger, Julie Han, Olivia B. Holmes, Yun-Fang Juan, Ryan King, Andrey Kislyuk, Maria Mariano, Lucia V. Reynoso, David Rissato Cruz, Jonathan Sheu, Jennifer Tang, James Wang, Mark A. Zhang, Emily Zhong, Vida Ahyong, Sreyngim Lay, Sophana Chea, Jennifer A. Bohl, Jessica E. Manning, Cristina M. Tato, Joseph L. DeRisi

## Abstract

**Background:** Metagenomic next generation sequencing (mNGS) has enabled the rapid, unbiased detection and identification of microbes without pathogen-specific reagents, culturing, or *a priori* knowledge of the microbial landscape. mNGS data analysis requires a series of computationally intensive processing steps to accurately determine the microbial composition of a sample. Existing mNGS data analysis tools typically require bioinformatics expertise and access to local server-class hardware resources. For many research laboratories, this presents an obstacle, especially in resource limited environments.

**Findings:** We present IDseq, an open source cloud-based metagenomics pipeline and service for global pathogen detection and monitoring (https://idseq.net). The IDseq Portal accepts raw mNGS data, performs host and quality filtration steps, then executes an assembly-based alignment pipeline which results in the assignment of reads and contigs to taxonomic categories. The taxonomic relative abundances are reported and visualized in an easy-to-use web application to facilitate data interpretation and hypothesis generation. Furthermore, IDseq supports environmental background model generation and automatic internal spike-in control recognition, providing statistics which are critical for data interpretation. IDseq was designed with the specific intent of detecting novel pathogens. Here, we benchmark novel virus detection capability using both synthetically evolved viral sequences, and real-world samples, including IDseq analysis of a nasopharyngeal swab sample acquired and processed locally in Cambodia from a tourist from Wuhan, China, infected with the recently emergent SARS-CoV-2.

**Conclusion:** The IDseq Portal reduces the barrier to entry for mNGS data analysis and enables bench scientists, clinicians, and bioinformaticians to gain insight from mNGS datasets for both known and novel pathogens.

## BACKGROUND

Infectious diseases remain a leading cause of morbidity and mortality worldwide. Despite significant advancement in our understanding of infectious disease biology, existing microbiological tests often fail to identify etiologic pathogens in cases of suspected infection. This can be due to a number of causes - failure to isolate an appropriate sample type, preemptive antibiotic exposure precluding growth in culture, lack of suspicion of a particular infection precluding the ordering of an appropriate test, or lack of available specific diagnostic tests due, in part, to limited knowledge of circulating pathogens. This is compounded further by the fact that novel, previously uncharacterized pathogens may also be present. This fact was illustrated vividly by the recent emergence of COVID-19 in Wuhan, China, in early December 2019. Metagenomic next-generation sequencing (mNGS) of nucleic acid from biological samples offers the potential for a universal pathogen detection method, including the detection of novel species. mNGS has great potential as a broad spectrum surveillance or patient monitoring tool, especially in low and middle income countries where the infectious disease burden remains high [1]. While the expense of sequencing continues to drop, the challenge of mNGS data analysis, the lack of bioinformatics expertise, and the access to sufficient compute and storage remains a major obstacle.

mNGS experiments result in millions of sequencing reads generated from the nucleic acid present within a biological sample, which may include complex microbial populations. A primary goal of mNGS data analysis is to determine what nucleic acid derives from the host (for example, a patient), and what cannot be attributed to the host or environmental contaminants. Further analysis of the non-host sequence may then attempt to determine the relative abundances of different taxa present in a particular sample, as this may provide insight into the presence and relevance of potentially pathogenic microbes. This is typically done via alignment of sequencing reads to a reference database. In the context of infectious diseases, identification of pathogens via this approach obviates the need for pathogen-specific reagents or the ability to culture the microbe. This is especially important for microbes that are difficult, or impossible to culture, including many viruses, fungal species, eukaryotic parasites, and bacteria [2]. Additional downstream analysis may then be employed to understand trends in the abundances and relatedness of organisms across samples.

There are several tools available for estimating relative abundance of microbial populations from mNGS data [3–20]. However, running these tools requires bioinformatics expertise and fluency with command-line tools. Additionally, pathogen detection in the context of a host organism presents unique informatics challenges beyond microbial abundance estimation. As noted, a substantial fraction of the sample may consist of host sequences that are secondary to the goal of pathogen detection [21]. Existing tools do not perform sensitive removal of host sequences or quality control (QC) steps, thus requiring the use of separate QC and alignment tools, and therefore additional computational experience in pipelining. A number of tools exist to incorporate multiple pipeline steps alongside reporting capabilities, including OneCodex [22], Sunbeam [23], and SURPI [24]. However, these tools require paid subscription or significant computational resources to build the underlying databases and run the analyses. Consequently, existing tools are not sufficient to support new applications of mNGS in poorly resourced settings where the detection of infectious agents could make a major impact on population health.

Here, we describe IDseq - an open source cloud-based service for pathogen detection and monitoring. IDseq is a continuously evolving service that enables robust and reproducible analysis of mNGS data for microbial identification, regardless of sample type or host species. We first describe the technical aspects of the IDseq pipeline implementation, including host filtration and QC, assembly-based alignment, and downstream reporting and visualization tools. We then evaluate the performance of the IDseq pipeline, first on a set of standard mNGS benchmark samples as compared to other tools aimed at providing taxonomic abundance estimates from mNGS data, and secondly on a simulated dataset to evaluate the ability to detect divergent viruses. Finally, we provide two case studies to demonstrate the application of IDseq. First, in a subset of samples from a previously published report which sought to investigate unknown etiologies of pediatric meningitis [1]. Secondly, we describe the performance of IDseq in the context of a real-world nasopharyngeal swab, processed and uploaded to IDseq from Phnom Penh, Cambodia, with respect to an emerging viral pathogen, SARS-CoV-2. By combining an intuitive web application, a cloud-based pipeline, and downstream visualization tools, IDseq enables investigation of mNGS data for pathogen detection and monitoring, especially suited for researchers with limited computational resources. Importantly, IDseq also enables facile collaboration and data sharing, while enhancing data analysis reproducibility across organizations and countries.

## IMPLEMENTATION

### IDseq Bioinformatics Pipeline

The IDseq Portal https://idseq.net is a cloud-based, open-source bioinformatics platform that enables detection of microbial pathogens from raw next-generation sequencing (NGS) reads. The IDseq pipeline is conceptually based on previously implemented pipelines [1,25], but is optimized for scalable Amazon Web Services (AWS) cloud deployment (**Figure 1**). Here, we describe v3.13 of the IDseq pipeline. Up-to-date pipeline documentation can be found at https://help.idseq.net.

**Figure 1:**
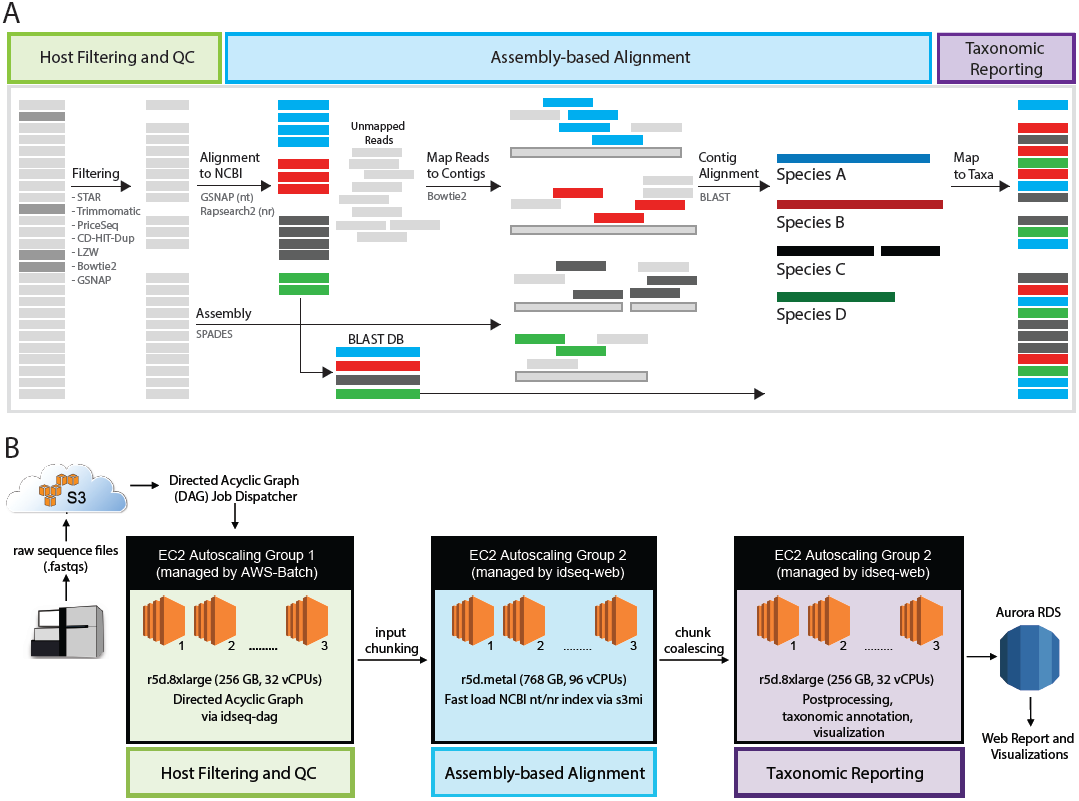
**A)** Overview of the IDseq pipeline steps and data analysis workflow. The IDseq pipeline for pathogen discovery is composed of several steps, including host filtering and QC, assembly-based alignment, and taxonomic aggregation and reporting. Each step is comprised of a number of existing bioinformatics tools. **B)** The IDseq pipeline is optimized for AWS cloud computational infrastructure. Each of the core pipeline steps (host filtering and qc, assembly-based alignment, and taxonomic aggregation and reporting) is managed by EC2 Autoscaling Groups.

The IDseq pipeline ingests raw, short-read sequencing data (either RNA- or DNA-seq from any sample type), which can be uploaded from local sources via the web interface or the command line interface (CLI) or directly from Illumina’s BaseSpace platform. Sequence analysis proceeds through three main phases: 1) Host filtering and QC, 2) Assembly-based alignment, 3) Reporting and visualization (**Figure 1A**).

#### Host Filtering and QC

The first phase of the pipeline begins with validation of input files (single- or paired-end .fastq or .fasta files). Currently, raw read files are arbitrarily capped at 150 million reads, a threshold that is, according to our experience, larger than most single metagenomic samples. Most mNGS samples processed for pathogen detection are sampled from a potentially infected host organism and thus the majority of sequencing reads derive from the host organism itself [21]. IDseq performs *a priori* subtraction of host sequences via STAR (Spliced Transcripts Alignment to a Reference) alignment of raw reads to a host-specific database [26]. IDseq is host-agnostic and allows researchers to select from several available hosts including human, mouse, pig, ticks, and mosquito, among others. For example, human host samples are aligned to the HG38 reference database (GCA_000001405.15), while mosquito samples are aligned to a combined collection of reference genomes from *Culex* and *Aedes* species as well as other diptera. Reads that align to the selected host genome are removed from the analysis. For hosts with well-annotated genomes, individual gene counts may be saved for offline transcriptome analysis, provided appropriate consent in the case of human subject research. Such host-based analyses have been shown to complement metagenomic analysis for pathogen detection [27]. For all host organisms, sequences for optional spike-in RNA controls developed by the External RNA Controls Consortium (ERCCs) are automatically recognized for downstream steps.

Next, IDseq performs a series of Quality Control (QC) steps, as outlined in **Figure 1**. First, Trimmomatic [28] trims Illumina adapters. Low-quality reads, duplicates, and low-complexity reads are then removed using the Paired-Read Iterative Contig Extension (PRICE) computational package [29], the CD-HIT-DUP tool (v 4.6.8) [30], and a filter based on the Lempel-Ziv-Welch (LZW) compression score, respectively. Regardless of the host genome, the data is scoured to remove all remaining human sequences using Bowtie2 against the HG38 reference database [31] and gmap-gsnap against a more stringent database including sequences combining both HG38 and Chimpanzees (*Pan troglodytes*) [32]. This step is especially important in the case of vector research, where blood meals may contain human sequences. At each step, the total number of reads remaining in the analysis is computed and these basic QC metrics (including total non-host reads, % passing QC, and duplicate compression ratio) are provided both in the user interface, as well as via download.

While the host filtering and QC steps performed by the IDseq pipeline serve primarily to reduce the computational burden and noise in downstream alignment steps, these metrics can also provide a resource for evaluating and troubleshooting sample preparation steps. The proportion of reads lost at each step may provide insight into possible sample degradation, fragment size, sequencing quality, or library complexity. IDseq’s automatic estimation of ERCC abundances enables back-calculation of the total input nucleic acid content, estimation of the lower limit of detection, and increases the ability to distinguish contaminants [34]. ERCC spike-ins are increasingly recognized as a best-practice for addressing the challenges associated with distinguishing background contamination from true microbial populations (Methods) [33].

#### Assembly-based Alignment

To assign taxonomic identities to each read, an assembly-based alignment procedure is used. First, filtered short-read sequences are aligned to the NCBI nucleotide (nt) and non-redundant protein (nr) databases (ftp://ftp.ncbi.nlm.nih.gov/blast/db/FASTA/) using GSNAPL [32] and RAPsearch2 [35], respectively (**Figure 1A**). GSNAPL is a specialized instance of the gmap-gsnap package written by Tom Wu, intended for very large genome databases.

The NCBI database indices are updated biannually, or as needed, via direct pull from NCBI. The index version is tracked for each pipeline run providing for versioned results. Putative accessions are assigned to each read using the NCBI accession2taxid database [36] and a BLAST+ (v 2.6.0) [37] database is constructed on-the-fly from the set of putative accessions (one database for each, nt and nr). In parallel, short reads are *de novo* assembled into contigs using SPADES [38]. Raw reads are mapped back to the resulting contigs using Bowtie2, in order to identify the contig to which each raw read belongs. Finally, each contig is aligned to the set of possible accessions represented by the BLAST database generated in the previous step, thereby improving the specificity of alignments to all the underlying reads, especially for homologous regions where short reads may align equally well to multiple different accessions.

### AWS Cloud Infrastructure

IDseq is optimized for scalable Amazon Web Services (AWS) cloud deployment (**Figure 1B**). Bioinformatics data processing jobs are orchestrated by the IDseq pipeline directed acyclic graph (DAG, https://github.com/chanzuckerberg/idseq-dag) and carried out on demand as Docker containers using AWS Batch. Alignments to the National Center for Biotechnology Information (NCBI) database are executed on dedicated auto scaling groups (ASG) of Amazon Elastic Compute Cloud (EC2) instances, with the number of server instances varied with job load. Fast downloads of the NCBI database from the Amazon Simple Storage Service to each new server instance are enabled by the open-source tool s3mi (https://github.com/chanzuckerberg/s3mi).

### Reporting and Visualization

Where alignments exist, taxonomic identifiers (taxID) for each of nt and nr, are assigned to each read. If there exist alignments with equivalent scores to multiple species taxIDs, then a single taxID is selected at random. If a read was incorporated into a contig, it is assigned the taxID belonging to the NCBI accession to whom its parent contig was assigned, as described above. If the read does not assemble into a contig, it is assigned the taxID of the NCBI nt and nr accessions it mapped to in the initial short-read alignment phase. The results are then aggregated to produce NT and NR counts for each taxID at both the species and genus level. Reads matching GenBank records in the superphylum Deuterostomia are removed, given the high likelihood that such residual reads are of host origin.

The IDseq Portal provides a number of different methods for interpretation of the pipeline results (**Figure 2**). First, relevant QC metrics and pipeline run information, including the number of reads remaining at each step of the host and quality filtering steps as well as estimates of internal control abundances are provided for each sample (**Figure 2AB, Methods**). The single-sample report tables provide key metrics for each taxon identified in the sample, including the total number of reads aligning to the taxon (in both NT and NR) as well as contig stats from the assembly-based alignment step (**Figure 2C**). The tree view enables rapid assessment of taxonomic relatedness of microbes identified in the sample (**Figure 2D**). For all views of the data, a wide range of user-selectable compound query and filtering tools are made available, enabling facile investigation of the data. For each taxonomic category, IDseq also provides one-click downloads of the corresponding underlying reads and contigs. Furthermore, coverage plots for contigs relative to all corresponding accessions to which they map are automatically generated (**Figure 2F**). To assist with distinguishing microbial signal from reagent and environmental contamination, IDseq supports background model generation, which allows researchers to evaluate the significance (reported in z-scores) of relative abundance estimates for taxons in samples of interest as compared to water-only or other environmental control sample collections. Altogether, the single sample report and associated filtering functionality enables evaluation of taxonomic hits. More documentation on specific metrics can be found at https://help.idseq.net.

**Figure 2:**
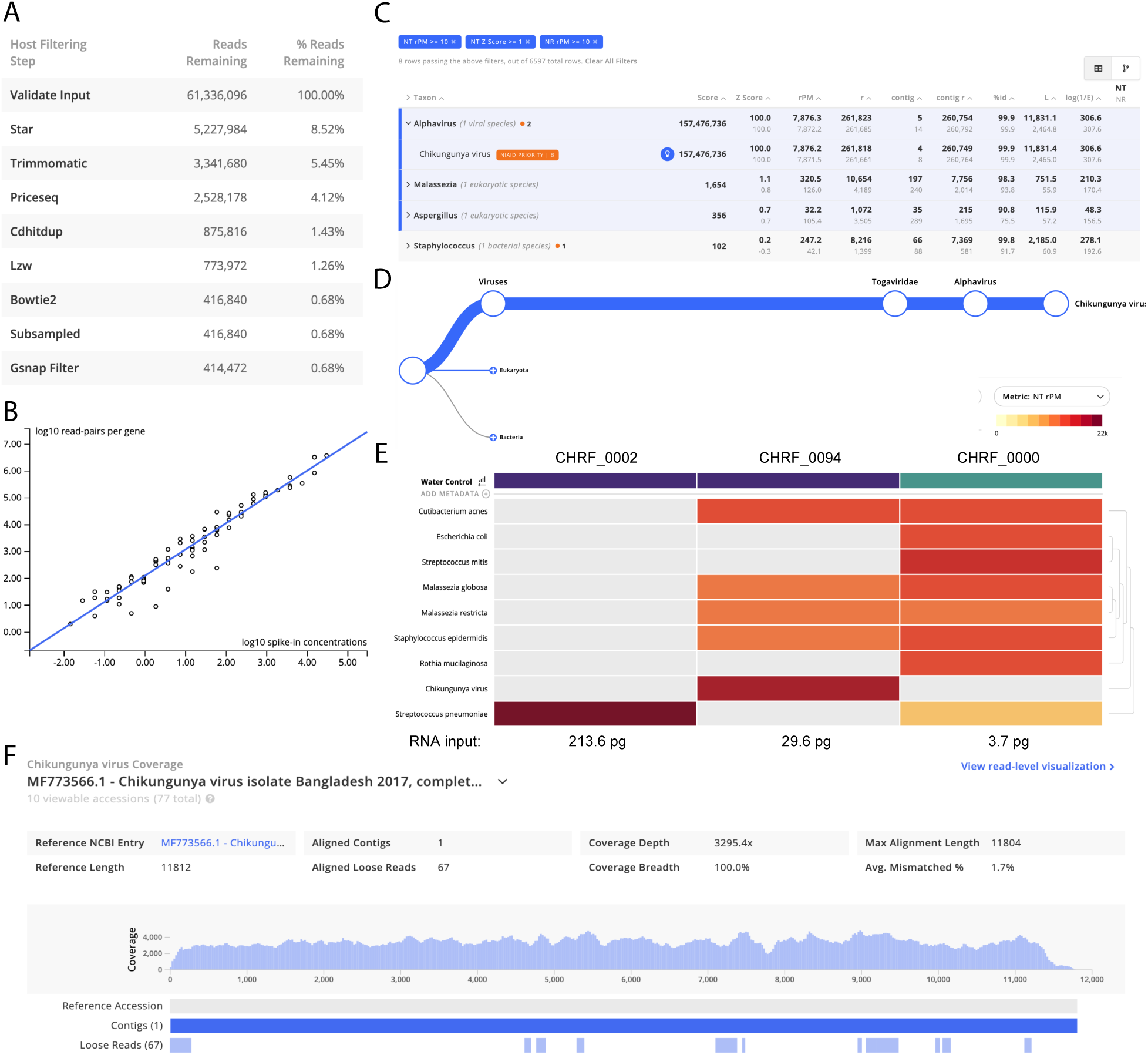
The IDseq web application provides multiple easy-to-use visualizations to help the user assess the quality and content of their sample. Screenshots taken from the IDseq Portal correspond to the re-analysis of samples from a study of etiologies of pediatric meningitis originally published in Saha *et al*. 2019 (see section: Application I). CHRF_0002 and CHRF_0094 are CSF samples from pediatric patients with meningitis due to *Streptococcus pneumonia* and *chikungunya virus*, respectively. CHRF_0000 is a water control. **A)** Table of reads remaining during each step of the host filtration step (for CHRF_0094) - interpretation of the relative loss at each step in can provide insight into the quality of the library preparation and sequencing run. **B)** Automatic quantification of ERCC counts from sample CHRF_0094; ERCC quantification enables back-calculation of input RNA concentration. **C)** The results for a single sample (CHRF_0094) are presented as a table, with key metrics for interpreting taxon alignment quality. **D)** The tree view indicates the relative abundance of sequences and their taxonomic relationship within a particular sample, shown is the relative abundance of *chikungunya virus* reads in CHRF_0094. **E)** The results from multiple samples can be compared using the IDseq heatmap view, with associated metadata (purple = CSF, blue = water control). The interactive heatmap visualization can be viewed at https://idseq.net/zlfl1. The heatmap is especially powerful when analyzing trends across a larger number of samples. **F)** Coverage of *chikungunya virus* in CHRF_0094; the coverage visualization enables rapid interrogation of genome coverage.

To facilitate visualization and hypothesis generation across multiple samples, the IDseq portal provides user customizable taxon heatmaps (**Figure 2E**). For advanced users, the pipeline visualization tool clearly documents the input parameters at each step of the analysis pipeline and provides download access to the input and output files at each step so data can be made available for offline analysis (**Figure S1**), such as phylogenetics.

### Versioning and Development

IDseq is an open source software tool under continued development across two GitHub repositories – one which hosts the web interface (https://github.com/chanzuckerberg/idseq-web), and one which hosts the pipeline code (https://github.com/chanzuckerberg/idseq-dag). Modifications to the web interface, which are deployed twice-weekly, do not affect the analysis results. To provide a record of features and how to use them, full documentation is provided at https://help.idseq.net.

Updates to the pipeline code may impact analysis results. Therefore, IDseq has adopted a semantic versioning system. Changes implemented to each version are listed in the README file. For each pipeline run, the pipeline and NCBI database versions are also tracked. Major changes to the pipeline outputs result in a major version number update (2.x to 3.x) and are communicated broadly to researchers via email updates. The change from IDseq v 2.x to v. 3.x involved the incorporation of the current assembly steps to refine alignment results, which improved the ability to resolve taxonomic identities in potentially homologous regions. Small changes to the pipeline that may still affect downstream results are indicated by an increased minor version number. For example, addition of a minimum alignment length filter to improve specificity of NT alignments caused a version change from 3.9.4 to 3.10.0. Changes to the pipeline which do not affect the results are indicated by incremental minor version (i.e. 3.13.1 to 3.13.2).

Continued development on IDseq aims to 1) improve the computational efficiency and accuracy of the results; 2) expand the integration with other tools to enable researchers’ flexibility in the downstream analysis of their processed results; 3) support the expanding number of mNGS sequencing platforms that will be used by researchers for pathogen detection globally. A suite of benchmarking samples are used for analysis of additional pipeline updates as discussed below.

### Software and Data Availability

Additional documentation and guides for getting started with IDseq can be found at https://help.idseq.net. The code is open source and available in the GitHub repositories listed in **Table S1**.

## RESULTS

### Evaluation of IDseq on External Benchmark Datasets

#### IDseq Analysis of Unambiguously Mapped Datasets

A recent study evaluated the performance of 20 taxonomic classifiers for mNGS data on ten reference datasets that are commonly used for benchmarking, containing computationally simulated reads from between 12 and 525 bacterial species [39]. It evaluated performance using two metrics - the area under the precision recall curve (AUPR) and the L2 distance. The AUPR evaluates the ability to detect the presence of microbes (binary presence/absence) above a relative abundance threshold, taking into consideration the precision and recall rates as said threshold is adjusted. A species-level AUPR of 1.0 indicates that there is a threshold (proportion of reads) above which all true positive species can be identified with no false positive species. The L2 distance provides a complementary metric that considers the similarity in relative abundances between the results and the ground truth.

We evaluated the performance of the IDseq pipeline on these same datasets (**Methods, Table S2**). Samples took an average of 3 hours (min = 1.6 hours, max = 10 hours) to process on IDseq pipeline version 3.13, with the NCBI database version from September 2019. Performance metrics (AUPR and L2 distance) were computed separately for the NCBI nt and nr results and compared to those published recently by Ye *et al*. (idseq_nt and idseq_nr, **Figure 3**) [39]. IDseq provides an automated pipeline, but at the cost of inability to easily swap in new databases. Therefore, we compared our results against those reported by Ye *et al*. for the “default database” of other tools. The performance metrics may inherit biases due to differences in the reference database contents as well as recency of input sequences.

**Figure 3:**
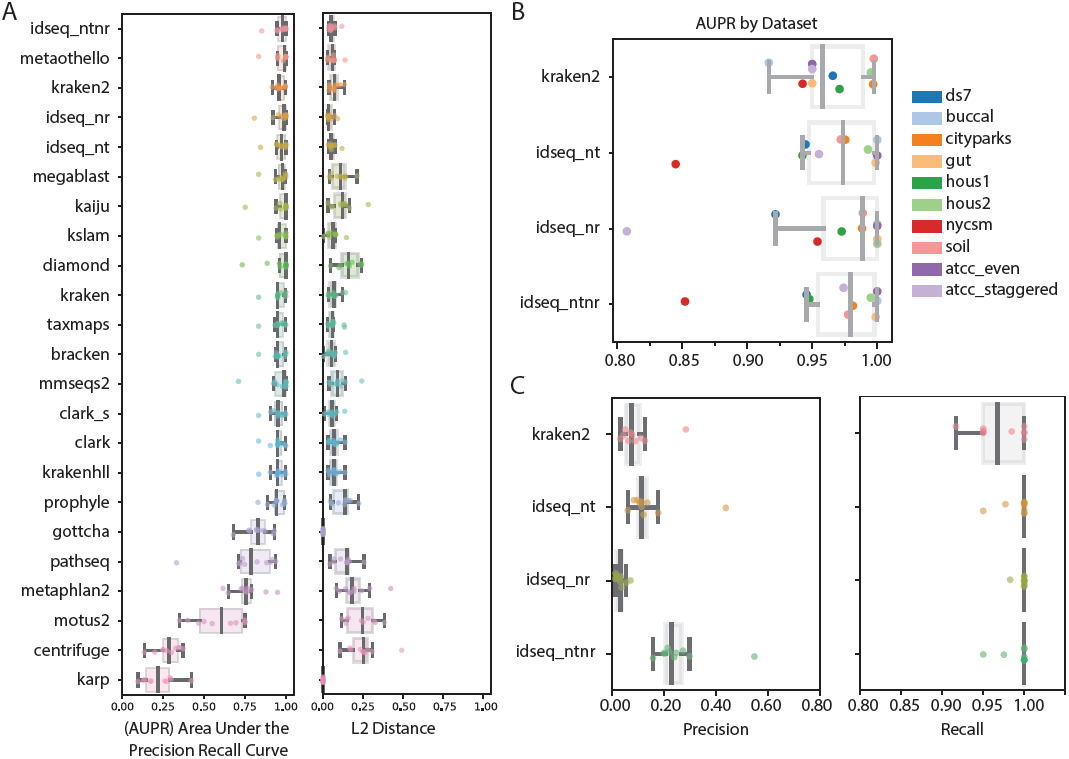
Performance metrics calculated for IDseq (NT and NR), as compared to the values recently published by *Ye et al*. [39] **A)** Area under the precision recall curve (AUPR) and L2 distance values for 22 tools, as evaluated against their default databases. **B)** The AUPR values for specific benchmark datasets evaluated for three tools (Kraken2, IDseq NT, and IDseq NR), including metrics obtained when evaluating basic threshold filters integrating both IDseq NT and NR (idseq_ntnr). **C)** The precision and recall of the same three tools for detecting known taxa.

#### Deep Dive of Unambiguously Mapped Datasets Results

The IDseq pipeline demonstrated comparable performance to the other mNGS tools tested (**Figure 3**). The unambiguously mapped datasets demonstrated limited resolution for distinguishing the tools when evaluated by AUPR and L2, as most tools show relatively high performance (with AUPR scores above 0.8 at the species level, **Figure 3A**). Consistent with Ye *et al*, we observed that the greatest differences between tools was in the reduced precision at high recall. IDseq protein alignments (NR) demonstrated greater AUPR than IDseq nucleotide (NT) across most datasets, but consistently identified more taxa at low abundance (less than 1%), therefore resulting in reduced precision (**Figure 3C**). Meanwhile, IDseq NT exhibited increased specificity. IDseq NT and NR had a mean AUPR across all the datasets of 0.9627 and 0.9633, respectively. The top mean AUPR of any single tool was achieved by metaothello (0.9661), followed by Kraken2 (0.9635). Given Kraken2’s performance on the unambiguous benchmark datasets and its wide adoption for relative microbial abundance estimation, additional analyses focused on comparison against Kraken2 (**Figure 3BC**). Another distinguishing factor between the tools was in the number of reads that were “unclassified” across multiple datasets. mmseq2, metaothello, kaiju, and bracken consistently left > 10% of reads “unclassified”. IDseq (NT and NR) removed an average of 10% of the reads during host filtering and QC steps, but of the remaining sequences, an average of less than 1% of reads were unmapped across the ten datasets. This can be attributed in part to IDseq always assigning reads to a species when an alignment exists (increasing sensitivity at the expense of specificity) and secondly to the use of assembled contigs to refine alignments where short reads may have been unmapped.

To further investigate differences between the tools, we evaluated the results for each dataset independently (**Figure 3B**). IDseq NR demonstrated lower precision across all datasets than many other tools, including IDseq NT and Kraken2 (**Figure 3C, Figure S2**). The ATCC Staggered dataset, which includes several microbes present at very low abundance, yields the lowest AUPR of all samples tested via IDseq NR, consistent with findings in Ye *et al*. that protein-based classifiers consistently struggled to identify the low-abundance taxa amongst other low-abundance false positives. Meanwhile, IDseq NT demonstrated reduced performance on the NYCSM dataset (Supplemental Text). IDseq’s usage of the full NCBI nt and nr databases resulted in relatively high performance for the Buccal dataset. Ye *et al*. discuss that the Buccal dataset was a low-performing outlier for most evaluated classifiers due to inclusion of reads from a species with only contig-quality reference, which is not included in most default databases.

The IDseq web portal is designed to provide researchers with the choice of utilizing either NT or NR results, or both in conjunction with each other. For example, the impact of spurious NR alignments can be mitigated by requiring a corresponding alignment with IDseq NT. Using this strategy, the performance of IDseq was evaluated, considering the NT relative abundances reported for taxa with both NT r > 0 and NR r > 0 (idseq_ntnr, **Figure 3, Figure S2**). We observed that requiring concordance resulted in the greatest mean AUPR across all other tested tools (0.9673) and increased the precision of IDseq above that of either NT or NR alone.

Altogether, these results highlight some key trade-offs with respect to relative abundance estimation of bacterial species. IDseq is capable of identifying organisms with respect to the latest versions of NCBI and demonstrates relatively high recall (**Figure 3D**). But use of the full NCBI database may result in false-positive alignments at low abundance which can reduce precision (**Figure 3C**). In the context of pathogen-identification, it has been observed that infecting agents may comprise the majority of sequencing reads in certain circumstances [27]. For such data sets, the reduced precision for abundance estimation at low levels is less impactful. Meanwhile, researchers interested in evaluating highly complex microbiome composition at the species- and strain-level may need to bring in other tools to supplement their analyses [40–42] or rely on genus-level estimates provided by IDseq.

### Evaluation of IDseq on Internal Benchmark Datasets

To address the gaps between the existing benchmark datasets and the IDseq pipeline’s primary use-case for pathogen detection, we tested IDseq’s performance on three additional datasets specifically designed to evaluate detection of divergent viruses (**Methods**) and common clinical microbes (**Supplemental Text**). For each dataset, we evaluated the performance of IDseq (NT and NR), as compared to Kraken2 [15], using per-species recall.

#### Detection of Divergent and Novel Viruses

Viruses are known to evolve rapidly and therefore their sequences may diverge from sequences in the known NCBI database over relatively short timescales [43]. Maintaining the ability to detect divergent viruses is of paramount concern, given their role in numerous recent outbreaks, including the recent emergence of SARS-CoV-2, the coronavirus responsible for the COVID-19 outbreak [44–47]. The idseq-bench tool was used to generate 17 simulated NGS samples from Rhinovirus C genomes at varying levels of divergence (after in-silico forward evolution from a reference sequence obtained from the NCBI database), ranging from 100% identical to the reference sequence to 25% similar (at the nucleotide level) (**Methods, Figure 4A, Table S3**). The resulting samples were uploaded to IDseq (Project HRhinoC Simulation). Meanwhile, the same raw .fastq files (prior to host filtering), were analyzed using Kraken2 (**Methods**).

**Figure 4:**
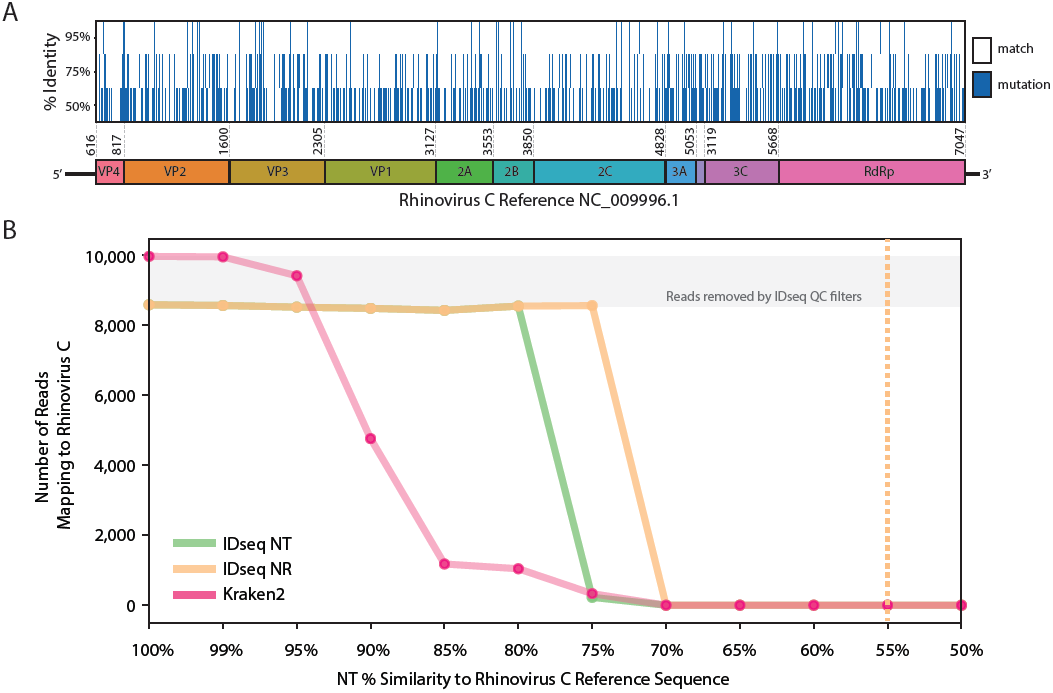
**A)** Graphic representation of genomic similarity for simulated divergent Rhinovirus C genomes, at 95%, 75%, and 50% similarity to reference sequence NC_009996.1. Mutations are shown in dark blue. **B)** Performance of IDseq (NT and NR) as compared to Kraken2 for recovery of reads from simulated divergent Rhinovirus C genomes at varying levels of divergence. The dotted yellow line indicates the theoretical limit for detection of Rhinovirus C achieved by manual BLASTx of IDseq-produced contigs.

Both IDseq NT and Kraken2 identified reads aligning to Rhinovirus C down to 75% sequence divergence (**Figure 4B**). Meanwhile, IDseq NR recalled Rhinovirus C alignments down to 70% sequence divergence, demonstrating a greater sensitivity for divergent virus detection. We note that IDseq NR experienced a rapid drop in total recall (8,558 reads correctly mapping to Rhinovirus C, of 10,000 total and 8,558 passing QC steps at 70% sequence similarity vs. 0 reads detected at 65% sequence similarity). This highlights an artifact of the computational cost-saving mechanisms employed by IDseq - whereby a BLAST database is constructed from only the subset of accessions identified in the initial short-read GSNAP and Rapsearch2 alignments to the NCBI database. In cases where the highly divergent short-read sequences don’t match to NT or NR in the initial alignment, the BLAST database will be empty and none of the reads or contigs will map. However, IDseq does provide the ability to download all assembled contigs, enabling offline interrogation of this divergent “dark matter”. Manual BLASTx of contigs assembled by SPADES in IDseq to the full NCBI database, was able to recover the Rhinovirus C identity down to 55% sequence identity. Future iterations of the IDseq pipeline may aim to automate the manual follow-up steps for divergent viral contigs as well as incorporating other tools for dark matter investigation to probe for pathogen motifs.

Further comparison of the IDseq (NT and NR) results to Kraken2 shows that Kraken2 initially recovered more of the simulated reads than IDseq (9,964 of 10,000 vs. 8,582 for both IDseq NT and NR). This is explained by the QC steps in the IDseq pipeline, which removed ∼15% of reads at the PriceSeq filtering step due to low quality - an expected outcome given that the simulated reads mimic error models of Illumina sequencers (**Methods**). Of the reads remaining after host filtering, IDseq identified 100% as aligning to Rhinovirus C. This pattern persists down to 95% sequence similarity, at which point Kraken2 begins to identify fewer reads. While some Rhinovirus C reads are identified down to 75% sequence similarity (same as IDseq NT), IDseq NT identified a significantly greater number of reads mapping to Rhinovirus C at increasing levels of divergence. Specifically, at 80% divergence, 8,544 reads mapped by IDseq NT while only 1,042 reads mapped by Kraken2. Altogether, these benchmark results are consistent with existing reports of the utility of IDseq NR in detecting divergent viruses [48] and are within the ranges of nucleotide divergence associated with emerging human pathogens (**Supplemental Text**).

### APPLICATION I. IDseq for Pathogen Discovery in Cases of Pediatric Meningitis

The IDseq pipeline is sample-type agnostic, allowing researchers interested in a broad range of scientific questions across a diverse array of host organisms (humans, mice, mosquitos, ticks, plants, environmental, etc.) to obtain relevant microbial information from any sample type (blood, CSF, respiratory fluids, tissue, etc.) [1,49–51]. There are many challenges for data interpretation that are common across mNGS applications, such as impact of PCR amplification on samples with low amounts of input RNA, background contamination, and genomic similarity between short regions of related organisms. Here, through a re-analysis of the IDseq results for three cerebrospinal fluid (CSF) samples from a recent study investigating etiologies of pediatric meningitis in Bangladesh [1], we highlight specific IDseq features to address these challenges. The original study, conducted by Saha *et al*. included 91 CSF samples (36 positive, 30 negative, and 25 idiopathic) and 6 water controls, processed on IDseq v3.1. We focus on one known infection (*Streptococcus pneumoniae*, CHRF_0002), one idiopathic sample that was later confirmed to have chikungunya virus (CHRF_0094), and one water control (CHRF_0000) (**Figure 2**). These samples, for demonstrative purposes, were re-run on IDseq v3.13 and are available in IDseq project CHRF RR007 Example. Key pipeline run metrics for these three samples are provided in **Table 1**.

**Table 1:** IDseq provides key metrics enabling The Host filtering and QC stage of the IDseq pipeline is composed of several individual steps. The proportion of reads lost at each step can provide insight into sample quality and library preparation. Interpretation of these metrics may be valuable for labs evaluating new sample storage techniques, library preparation protocols, etc. These three samples, provided as an example, can be investigated in the IDseq portal in Project <u>CHRF RR007 Example</u>. CSF: Cerebral Spinal Fluid, DCR: duplicate compression ratio, “number of rows”: total number of species and genus-level rows in the IDseq sample report, NT: results based on NCBI nucleotide (nt) database, NR: results based on NCBI non-redundant protein (nr) database, rPM: reads per million, L: average alignment length across all reads and contigs mapping to that taxon.

| Sample ID | CHRF_0094 | CHRF_0002 | CHRF_0000 |
| --- | --- | --- | --- |
| Description | Chikungunya viral meningitis | Streptococcus pneumonia meningitis | Water Control |
| Sample Type | CSF | CSF | H2O |
| Collection Location | Bangladesh | Bangladesh | Bangladesh |
| Total Reads | 61,336,096 | 141,979,356 | 135,087,088 |
| ERCC Reads | 28,094,424 | 14,875,054 | 130,150,782 |
| STAR | 5,227,984 | 26,802,224 | 4,675,916 |
| Trimmomatic | 3,341,680 | 23,770,970 | 3,440,016 |
| PRICE | 2,528,178 | 20,752,710 | 1,964,846 |
| DCR | 2.89 | 1.39 | 4.67 |
| RNA input concentration<br>(Back-calculated from ERCCs) | 29.6 pg | 213.6 pg | 3.7 pg |
| Number of rows<br>(NT rpm > 10, NR rpm > 10, NT Z > 1) | 8 | 2 | 36 |
| NT rPM | 7,876.2 | 22,034.0 | NA |
| NR rPM | 7,871.5 | 19,743.9 | NA |
| Number of Contigs | 4 | 143 | NA |
| Alignment L (nt) | 11,831.4 | 75,065.3 | NA |
| Average % Identity | 99.9 | 99.2 | NA |

#### Sample 0094: A case of neuroinvasive chikungunya virus

CHRF_0094 was a pediatric encephalitis case of unknown etiology that was later determined to be a case of neuroinvasive chikungunya virus. **Figure 2A** shows the number of reads removed by each host filtration and QC step. One challenge for mNGS-based pathogen detection is that host sequences dominate the mNGS library. Notably, in CHRF_0094, chikungunya virus reads in sample CHRF_0094 represented less than 1% of the total sequencing reads. However, after IDseq’s host filtering and QC steps, it represented 63% of the remaining non-host reads. A second, widely acknowledged challenge for mNGS data interpretation is the presence of environmental contaminants. Best-practices suggest including at least one water control with every sequencing experiment [34,52]. To assist with interpretation of results with respect to control samples, IDseq implements a z-score approach (**Methods**) first described in Wilson *et al*. [53]. Z-score statistics computed by IDseq indicate the significance of relative abundance estimates in a sample as compared to the user-selected background controls - which may include water controls or healthy control samples. Z-score thresholds can be imposed to remove taxa that are prevalent in the water or healthy controls. In sample CHRF_0094, 8 rows (4 species from 4 genera) were reported with NT reads per million (rPM) greater than 10, NR rPM > 10, and z-score > 1 (a relatively stringent threshold employed to remove many of the low-abundance taxa for first-pass evaluation). 7,876.2 rPM were associated with chikungunya virus, of which many were associated with the 4 contigs aligning to chikungunya virus. By using the IDseq portal coverage visualization, which displays reads and contigs in association with their top matched GenBank accession, we observe that the longest contig, approximately 11kb, represented full-genome coverage of the nearest GenBank accession (**Figure 2F**).

#### Sample 0002: A case of known Streptococcus pneumoniae meningitis

In sample CHRF_0002, IDseq associated 1,927,505 reads by NT with the independently verified pathogen, *Streptococcus pneumoniae*, of which 98.1% were assembled into 143 contigs (**Table 1**). The average alignment length across all contigs and reads was 75,289.8 bp - driven largely by alignment of long contigs. Despite the large number of contigs and long alignment lengths, the GenBank accession with the greatest coverage (1.8mb LR216026.1 Streptococcus pneumoniae strain 2245STDY5775485 genome assembly, chromosome: 1) had 87.3% coverage breadth. This exemplifies a frequently observed pattern (which is even more pronounced in lower-coverage samples) - whereby coverage of larger bacterial genomes is lower than virtual genomes even for samples with a high proportion of mNGS reads associated with a particular microbe. For many cases, low coverage from mNGS data can preclude confident strain identification in bacterial species that may be useful in a clinical context. Furthermore, low coverage of the transcriptome (via RNA mNGS) may produce a large proportion of alignments in conserved rRNA regions which may be challenging to disambiguate.

#### Sample 0000: A water control

In sample CHRF_0000, the duplicate compression ratio (DCR) of 4.67 indicates the possibility of over-amplification of low biomass nucleic acid input (**Table 1**). This is common for water samples where low input nucleic acid is expected. The use of ERCC controls in the library preparation of these samples enabled back-calculation of the total input RNA concentration. This sample was determined to have 3.7 pg of total input RNA, while the two infected samples (CHRF_0094 and CHRF_0002) had 29.6pg and 213.6 pg, respectively. Thus, while the relative abundance values appear comparable to those in the infected sample, they represent significantly smaller quantities of raw nucleic acid (**Figure 2E**). In the original study all water and non-infectious controls (which had low white cell counts and therefore little host or pathogen nucleic acid) had input RNA quantities < 4 pg, enabling the use of an input nucleic acid threshold for inclusion in downstream analyses. Additionally, the top four organisms (by NT rPM) include *Providencia, Cutibacterium, Streptococcus*, and *Escherichia* - many of which are known environmental contaminants [33,54,55]. The 36 total rows (with NT rPM > 10, NR rPM > 10, and NT z-score > 1) are all present at relatively similar and low abundance levels, characteristic of background contaminants [33,34].

### APPLICATION II. Real-time Detection of Novel Coronavirus

IDseq is a globally accessible pipeline for mNGS analysis that has been shown through simulation and practice to be effective in identifying novel and divergent viruses. As an additional real-world example of this utility, we provide a vignette from the recent SARS-CoV-2 coronavirus outbreak. On January 30, 2020, a team of researchers from the CNM-NIAID (National Center for Parasitology, Entomology, and Malaria Control - National Institute of Allergy and Infectious Disease) collaboration in Cambodia obtained a nasopharyngeal swab sample from a patient with PCR-confirmed SARS-CoV-2 infection. The library preparation and sequencing were completed in-country by February 1, 2020 [56]. Analysis of the sample (4.5 million single reads) using IDseq against an NCBI database version from 2019-09-17, which did not contain the known sequences for SARS-CoV-2 that have since been deposited on NCBI, identified 571 reads aligned to the genus *Betacoronvirus*, with an average amino acid percent identity of 92.3% (by NR). The sample took 14 minutes to analyze end-to-end and the most abundant species was severe acute respiratory syndrome-related coronavirus, with 542 NT reads (22 contigs) and 571 NR reads (24 contigs), representing ∼33% genome coverage. To quantify the IDseq pipeline’s recall for SARS-CoV-2 sequences, we built a BLAST database from the 54 sequences associated with SARS-CoV-2, which had been deposited in NCBI between January and February 2, 2020 as a result of widespread efforts by the global science community. By BLASTing all non-host reads from the sample against the known SARS-CoV-2 sequence database, we identified 584 reads mapping to SARS-CoV-2. As compared to this ground-truth value, IDseq demonstrated 97.8% read-level recall. This indicates that for an emerging threat, IDseq was able to successfully provide information on the presence of a pathogen prior to the existence of full reference genomes associated with the organism. This identification was of paramount public health importance given unclear diagnostic accuracy in the beginning pre-pandemic state.

## DISCUSSION

We have introduced IDseq, an open source cloud-based pipeline and analysis service for metagenomic pathogen detection and monitoring. We described the pipeline analysis steps and demonstrated that the IDseq pipeline achieves comparable performance for taxonomic identification and relative abundance estimation as other tools in the field. We showed that IDseq is uniquely suited for detection of divergent viruses and has high sensitivity for detecting human pathogens. Finally, we have shown through two case-studies, how the IDseq portal enables researchers to rapidly generate insights into their samples’ quality, microbial content, and cohort trends. We further highlighted its real-time utility by describing how IDseq was used to analyze sequences associated with the emerging coronavirus SARS-Cov-2 prior to deposition of SARS-CoV-2 sequences into public data repositories. The IDseq web portal provides an easy-to-use access point for computationally intensive analysis of mNGS data. Its sample-type agnostic implementation enables its application for a broad range of research questions related to understanding distribution of microbes in a sample. The IDseq pipeline has been a key component in recent studies to understand undiagnosed causes of infection and survey the landscape of circulating pathogens, both in humans and animals [57,58].

Benchmarking of mNGS tools is a well-recognized challenge within the field [39,59,60]. The choices of tools, parameters, databases, and datasets may all influence the conclusions. Our aim in this study was simply to test performance relative to other tools. We compare IDseq’s default database (NCBI nt and nr) against the default databases for all other tools included by Ye *et. al*. Though it is possible that other tools’ performance would improve given a comparably large database, configuring these details requires computational expertise that directly opposes the readily usable nature of IDseq. IDseq continues to use the full NCBI nt and nr database given their advantages for detecting divergent viruses and incorporating data on novel bacterial pathogens. However, the large database size results in longer run-times and the lack of curation induces the potential for noise in alignment results due to errant sequence assignment errors upon upload to the NCBI databases. There is ongoing work by many researchers to evaluate curated databases for mNGS analyses, but for now IDseq continues to update its database biannually. To support continued benchmarking of IDseq and empower researchers to test IDseq’s performance for their particular applications, we have released the open-source idseq-bench tool, which was used to generate the divergent virus dataset and for evaluating the per-read recall results.

Beyond the informatics nuances between tools, IDseq provides clear advantages for researchers new to mNGS and computational data analysis. First, IDseq is designed and maintained by a team of engineers and managed as a software-as-a-service product, where user support is a key component. User support enables researchers to have confidence that they will obtain results in a timely fashion. Secondly, the tool’s user interface provides a series of advantages for users with limited computational expertise by reducing the challenges associated with installation and configuration as well as providing meaningful metrics for quality control and interpretation. It maintains transparency on individual pipeline steps through documentation (https://help.idseq.net), the pipeline visualization tool (**Figure S2**), and availability of downloads from intermediate files. Together, these resources help researchers new to mNGS get started quickly, while also providing tools to enhance skills in computational biology. Thirdly, the pipeline provides assurance of computational reproducibility, which is an increasingly appreciated priority within the scientific community as dataset sizes and analytical complexity increase. Lastly, the web-based user interface provides an access point for collaboration and networking – enabling researchers to collaborate seamlessly across countries and institutions, thereby building global networks of expertise that can be accessed by those in resource-scarce settings.

Finally, we highlight that IDseq is not a clinical tool and intended for research-only purposes. IDseq aims to be a valuable resource for researchers in the infectious diseases field but does not intend to become clinically validated. While IDseq can yield insights that inform public health policies, laboratory testing priorities, and real-time decisions for confirmatory clinical testing, clinical validation of the pipeline requires locking of the system for adherence to strict guidelines. IDseq will remain under continued development in order to 1) improve the computational efficiency and accuracy of the results; 2) expand the integration with other tools to enable researchers’ flexibility in the downstream analysis of their processed results; and 3) support the expanding number of mNGS sequencing platforms that will be used by researchers for pathogen detection globally. Some possible future directions for improvements to the IDseq pipeline have been discussed throughout. Notably, IDseq’s current assembly-based alignment steps results in failure to automatically identify divergent viruses beyond 70% divergent, while BLASTx of IDseq-generated contigs can enable detection down to 55% divergent. Automating full NCBI BLASTx of putative viral contigs would simplify offline analyses. Similarly, we showed that IDseq NR had reduced precision, which made relative abundance estimation of low-abundance taxa challenging. Allowing for non-species-specific mappings or propagating estimates of species-level ambiguity to increase species-level resolution for low-abundance taxa may provide another avenue for continued development. Finally, continued integration with other analysis tools and sequencing technologies will further enhance the usability of IDseq for mNGS data analysis.

IDseq reduces the need for much of the computational expertise and access to large-scale computing resources that have traditionally been barriers for conducting mNGS data analysis. The IDseq portal provides an easy-to-use interface that enables researchers around the world to upload samples and generate hypotheses with relevant implications for global health and infectious disease tracking as diseases emerge.

## METHODS

### Raw Pipeline Commands

The IDseq pipeline uses several publicly available academic bioinformatics tools. The raw commands and parameters used for each step in the pipeline are available for each pipeline version in the pipeline visualization (**Figure S1**), which can be viewed for any sample in IDseq. Technical documentation is available here: https://github.com/chanzuckerberg/idseq-dag/wiki.

### Automatic ERCC Quantification

The External RNA Controls Consortium (ERCC) developed a common set of external RNA controls that can be used to control for a variety of sources of variation on RNA expression attributed to experimental factors (including the quality of the starting material, the level of cellularity and RNA yield, the sequencing platform, and the person performing the experiment). In the context of pathogen detection, mNGS libraries often contain extremely low quantities of RNA input. It has been shown that during library preparation, samples with low input experience amplification background contaminants [34]. ERCC controls can be used to mitigate the effect of low input libraries and to quantify the total input. To enable researchers to rapidly assess the quality of their libraries and the limit of detection, IDseq provides ERCC counts for each sample. During the host filtering steps, the raw sequencing reads are aligned to the ERCC reference sequences and counts are generated by STAR –genecounts option [26]. These values are then available for download, as well as visualized in the user interface (**Figure 2B**).

### IDseq Z-score and Aggregate Score Metrics

Given the sensitivity of mNGS, it is common to identify contaminating microbial sequences derived from laboratory contaminants, reagents, collection tubes, etc. There exist numerous approaches to assist in distinguishing background contaminants from true microbes [34,53,61]. IDseq implements a previously described z-score method for background correction [53]. Researchers can create a background model by selecting control samples sequenced via their standard laboratory protocols or select from a default set of publicly available water controls. From the selected set of samples, the distribution of reads for each taxon is computed. The z-score field of the IDseq sample report is calculated as the z-score for each taxonomic ID based on its prevalence in the selected background model. If a particular taxonomic ID is not found in the set of control samples, then the z-score will be set to 100. If the taxonomic ID is not found in the sample, the z-score will be set to −100. The z-score metric also feeds into the “aggregate score”, which combines information from NT rPM, NR rPM, NT z-score, and NR z-score to provide an estimate of “microbial importance” for a particular sample based on the relative abundance both with the sample as well as in the background. This experimental metric aims to rank rare organisms that may be implicated in an infection higher, even if they are present only at low abundance.

### External Benchmarks - Datasets and Metrics

Datasets evaluated by Ye *et al*. in their benchmark analysis of 20 mNGS tools were downloaded from <gs://metax-bakeoff-2019>. The raw .fastq files were uploaded to IDseq (**Table S2**). The truth files for each of the datasets were obtained from https://github.com/yesimon/metax_bakeoff_2019 and are available in the *Notes* field of the IDseq metadata. The code developed by Ye *et al*. was downloaded from the GitHub repository. IDseq sample reports were downloaded upon completion and processed to produce species-level relative abundance estimates for each sample – specifically, the proportion of total reads (by NT and NR) was computed and used as input to the script. The IDseq results were processed in parallel with the data analyzed for the Ye *et al*. paper. The scripts used to run this analysis are available here https://github.com/katrinakalantar/idseq-benchmark-manuscript. Modifications to the original script are annotated as “##IDseq EDIT”. The computed metrics (AUPR, L2 distance, precision, recall, and f1-score) were then output as .csv files and plotted (**Figure 3, Figure S3**).

### idseq-bench: IDseq Benchmarking Tool

The idseq-bench tool (https://github.com/chanzuckerberg/idseq-bench) was developed as a resource to enable the IDseq team to benchmark datasets internally [62]. The tool is open source and available for external users to generate benchmarks appropriate for their particular use case. Full documentation can be found on GitHub. Briefly, the tool enables users to simulate NGS sequencing data from known microbes. By indicating the GenBank reference accession, idseq-bench uses the InSilicoSeq simulation tool [63] to generate reads in accordance with known sequencing error models. The true organism from which each read was simulated contains a tag indicating the known accession and species-, genus-, and family-level taxonomic IDs. The idseq-bench tool then uses this information to characterize performance of the IDseq pipeline results. The tool provides metrics for read-level-recall at the species-, genus-, and family-level, as well as sample-level AUPR, L2, precision, recall, etc. For samples that were not simulated internally, the tool enables users to supply a gold standard file (comparable to those obtained for the Cell Benchmarks Datasets) and compute sample-level metrics against that file.

### Internal Benchmarks - Divergent Virus Simulation and Analysis

A reference genome for Rhinovirus C (RefSeq NC_009996.1) was identified and the associated coding sequence .fasta file was downloaded from RefSeq. VIRAPOPS forward viral simulation [64] was used to simulate 5000 generations of viral evolution using default parameters. From the simulated data, sequences were selected at intervals of 5% nucleotide sequence identity to the original reference and compiled into a fasta file. This was then used as input to the idseq-bench simulation tool for benchmark simulation, which used InSilicoSeq [63] to simulate 10,000 sequencing reads of length 126 for each divergent virus genome according to a HiSeq error model. This resulted in 195.8x coverage of each divergent viral genome, consistent with the relatively high coverage of viral genomes seen by IDseq analysis of samples with high viral load. The simulated fastq files were then uploaded to IDseq project HRhinoC Simulation (Samples HRC_100, HRC_99, HRC_95, … HRC_025, **Table S3**).

To evaluate the limit of detection for divergent viruses, the total recall of Rhinovirus C reads was evaluated at each level of simulated divergence, for each tool. Additionally, the number of reads aligning to false-positive species was tracked. Offline analysis was done using the contigs generated by IDseq for samples where IDseq failed to identify Rhinovirus C. For simulated samples HRC_070 through HRC_025, the “unmapped contigs” were downloaded and aligned via BLASTx in the NCBI BLAST web interface using default parameters [37]. Samples for which the BLASTx result returned Rhinovirus C were marked as “potentially possible” and the greatest level of divergence was recorded.

### Internal Benchmarks - Running Kraken2

To compare internal benchmark samples against Kraken2 [15], a Kraken2 database was generated from the NCBI NT sequence database [65]. The following command line parameters were used to download and build the reference database. Finally, simulated sequencing files were run via the following commands.

Download the NCBI Database:

kraken2-build --download-library nt --db db_ncbi_nt

Build the Kraken2 NCBI Database:

kraken2-build --build --db db_ncbi_nt --threads 8

Run Kraken2 on benchmark datasets:

\# classify: running kraken changes slightly based on the sample being

compressed/decompressed or single/double pair

BENCHMARK=<benchmark_name_minus-R#> FORMAT=fastq bash -c ‘/usr/local/sbin/kraken2

--db databases/kraken2/ncbi_nt --threads 8 --gzip-compressed --classified-out

results/kraken2/$BENCHMARK.classified_seqs#.fq --unclassified-out

results/kraken2/$BENCHMARK.unclassified_seqs#.fq –output

results/kraken2/$BENCHMARK.kraken2.out –paired

benchmarks/${BENCHMARK}_R1.$FORMAT.gz

benchmarks/${BENCHMARK}_R2.$FORMAT.gz >

### Application I - Data Processing

In collaboration with Saha *et al*. [1], three samples were identified (CHRF_0000, CHRF_0094, CHRF_0002, from the original NCBI Sequence Read Archive dataset under BioProject PRJNA516582) and re-run on pipeline version 3.13. The pipeline results were filtered using a conservative set of filters, which required NT_rPM > 10 and NT_zscore > 1. The z-score was computed with respect to the public background model CHRF_RNA_Negative, which was used in the original manuscript. The background model was generated based on RNA-seq data from water samples and negative controls. Metrics were compiled into **Table 1** and a heatmap was generated using IDseq, with the same filters (**Figure 2D**).

### Application II - Data Processing

In collaboration with Manning *et al*. [56], RNA was extracted from a sample obtained from a symptomatic patient meeting criteria for possible COVID-19 pneumonia. Libraries were prepared for sequencing as described in Manning *et al* and sequenced on an Illumina iSeq100. The raw .fastq files were uploaded to IDseq from the CNM-NIH lab in Phnom Penh, via Illumina BaseSpace, on January 31, 2020 using an NCBI index from September 2019. An NCBI database update was then done on February 2, 2020 by the IDseq team and the results were evaluated. These samples were run on IDseq pipeline version 3.18. The data was deposited in public repositories by the original authors and is available at GISAID accession EPI_ISL_411902. IDseq results for the associated samples are available at http://public.idseq.net/covid-19?utm_source=bioarxiv&utm_medium=paper&utm_campaign=benchmark-paper.

## Availability of supporting source code and requirements

Project name: IDseq Portal

Project home page: https://idseq.net

Operating system(s): Platform independent

Programming language: Python, Ruby, JavaScript

Other requirements: Web browser

License: MIT License

## Availability of supporting data

Data referenced in this manuscript has been previously published. SRA accession IDs are included in the original manuscripts [1, 39, 56].

## Supplemental Text

Supplemental_Text.docx, contains supplemental methods and results associated with two benchmark datasets listed in the main text, as well as supplemental figures (Figure S1 – S3) and supplemental tables indicating IDseq data availability (Table S1 – S3).

## List of Abbreviations

ASG: Auto-scaling Groups
SARS-CoV-2: Severe Acute Respiratory Syndrome Coronavirus 2
AUPR: Area Under the Precision Recall Curve
AWS: Amazon Web Services
CLI: Command Line Interface
CSF: Cerebrospinal Fluid
DAG: Directed Acyclic Graph
DCR: Duplicate Compression Ratio
EC2: Elastic Compute Cloud
ERCCs: Externa RNA Controls Consortium
mNGS: metagenomic Next-Generation Sequencing
NCBI: National Center for Biotechnology Information
QC: Quality Control
taxID: Taxonomic identifier
NT: Nucleotide, NCBI nucleotide (nt) database
NR: Protein, NCBI non-redundant protein (nr) database
rPM: Reads per Million Reads Sequenced

## Acknowledgements

This research was supported by the Chan Zuckerberg Initiative (CZI). The authors would like to thank all CZI team members who were involved in support for software development and all researchers who have provided input on the IDseq Web Portal throughout the course of its development. The authors would like to thank Hanna Retallack for valuable comments on the manuscript text.

## Author’s Contributions

KLK conceived the project. KLK and JLD structured the draft and provided final editing. KLK coordinated and drafted the manuscript, and synthesized comments provided by all authors. All authors contributed critically important comments. VA, SL, SC, JAB, and JEM contributed to the generation of COVID-19 sequencing results. The IDseq Engineering Team (TC, CDB, BD, GD, RE, JH, OH, YJ, RK, AK, MM, LR, DRC, JS, JT, JW, MZ, EZ) contributed to software development. All authors read and approved the final manuscript.

## SUPPLEMENTAL TEXT

### Supplemental Methods

#### Internal Benchmark Datasets and Analysis

Many closely related bacterial species have different relative pathogenicity in humans or other host organisms [1]. The unambiguous benchmark datasets, evaluated in the main text, do not emphasize performance at distinguishing common clinically relevant pathogens and closely related bacterial species. Here, we provide the results of an analysis of two simulated datasets containing common clinical microbes. For each dataset, we evaluated the performance of IDseq (NT and NR), as compared to Kraken2 [2], using per-species recall.

#### Internal Benchmarks - Common Clinical Microbes (CCM) Benchmark Dataset

Several microbes commonly identified in samples from humans and vector samples were identified in collaboration with researchers and incorporated into the CCM benchmark dataset. The selected species represent a range of bacterial, viral, fungal, and parasitic pathogens. The idseq-bench simulation tool was used to pull reference genomes from 35 specified accessions across 13 species (*Species name*, taxonomic ID; *Klebsiella pneumoniae*, 573; *Aspergillus fumigatus* Af293, 746128; *Plasmodium falciparum*, 5833; Rubella virus, 11041; Human immunodeficiency virus 1, 11676; Rhinovirus C, 463676; Chikungunya virus, 37124; *Staphylococcus aureus*, 1280; *Balamuthia mandrillaris*, 66527; *Elizabethkingia anophelis*, 1117645; *Neisseria meningitidis*, 487; Torque teno midi virus, 2065052; Hubei mosquito virus 2, 1922926) and simulate reads using the InSilicoSeq [3] sequence simulator with uniform coverage and sequencing error models derived from HiSeq data. The simulated sample is available on IDseq (**Table S3**, https://idseq.net/ak8qu). This simulated dataset is utilized by the IDseq team to evaluate the consistency of IDseq results over time and ensure the reliability of the pipeline results.

For IDseq NT and NR results, the idseq-bench score function was used to evaluate the per-read, species-level recall across each ground truth organism. Then, to characterize false positives, the total number of species identified was determined, along with the percentage of reads mapping to incorrect taxa. For Kraken2, the read-level recall was determined at the species level and reads mapping to higher taxonomic levels were not considered as positive matches for the species-level recall. Again, the total number of unique species was determined and the proportion of reads mapping incorrectly at the species level was determined.

#### Internal Benchmarks - Closely Related Bacteria (CRB) Benchmark Dataset

Many bacterial pathogens are closely related, therefore resulting in possible genomic overlap and ambiguity in aligning short read sequences that may map to multiple species. However, distinction of pathogens at the species-level has implications for treatment and outcomes. 10 bacterial species from 6 genera within the Enterobacteriaciae family were identified and the idseq-bench simulation tool was used to pull reference genomes (*Species name*, taxonomic ID; *Salmonella enterica*, 28901; *Citrobacter koseri*, 545; *Citrobacter freundii*, 546; *Klebsiella aerogenes*, 548; *Enterobacter cloacae*, 550; *Escherichia coli*, 562; *Klebsiella oxytoca*, 571; *Klebsiella pneumoniae*, 573; *Shigella boydii*, 621; *Shigella flexneri*, 623) and simulate reads using the InSilicoSeq [3] sequence simulator with uniform coverage and sequencing error models derived from HiSeq data. The simulated sample is available on IDseq (**Table S3**, https://idseq.net/71wns). Analysis of the CRB benchmark dataset replicated that which was done for the CCM dataset (above).

### Supplemental Results

#### External Benchmarks - NYCSM Outlier Dataset and IDseq NT

The NYCSM dataset had the lowest AUPR of all datasets tested by IDseq NT. Overall, we note that due to the relative comparable performance of tools on these particular datasets, distinctions in AUPR may be the result of just one or two missed taxa. In this case, IDseq NT failed to identify *Enterobacter asburiae* (taxID = 61645) and identified only a small number (< 10) of reads aligning to *Pseudoalteromonas haloplanktis* (taxID = 228). Meanwhile, IDseq NR identified successfully recovered reads from *E. asburiae* and produced a comparably low number of reads mapping to *P. haloplanktis*. IDseq NT did identify many reads mapping to *Enterobacter soli*, which has been noted to have high genomic similarity to *E. asburiae* and *E. aerogenes* (p-distance: 1.06 and 1.19%, respectively [4]), both of which were included in the simulated dataset. This highlights the challenge with disambiguating short reads from taxa with a high degree of genomic similarity. We emphasize the practical importance of orthogonal validation of hits via assays (ie PCR) targeting unique regions of the genome. However, it is possible that the expansion of reference databases has improved specificity beyond the original dataset simulation. The particular GenBank accession to which all *E. soli* reads map (CP003026.1 Enterobacter soli strain LF7a, complete genome), was added to the database in April 2019, while reads were simulated in 2016 [5]. The same trend is observed for *P. haloplanktis*. IDseq NT and NR both identified significantly more reads mapping to *P. arctica*, a member of the *P. haloplanktis*-like group [6]. The associated GenBank accession to which the majority of reads map (CP011025.1 - Pseudoalteromonas arctica A 37-1-2 chromosome I, complete sequence) was added in September of 2017. The IDseq coverage visualization makes interrogation of microbial hits simple by linking to the NCBI taxonomy database and GenBank accessions.

#### Viral Divergence in Recent Human Disease

The ability to detect divergent viruses is a function of genomic similarity to other organisms in the database as well as genomic coverage, which influences assembled contig lengths. Many recent, emerging, diseases affecting humans do have some sequence similarity to organisms in the NCBI databases. For example, since the emergence of enterovirus EV-D68, numerous outbreaks caused by divergent sub-clades have been reported with nt similarity to other strains ∼96% [7]. Similarly, recent outbreaks of Dengue virus have been reported to be the result of introduction of novel DENV lineages, which are defined based on nucleotide divergence of 6-8% within each DENV serotype[8,9]. The set of nine known West Nile Virus lineages (which genomic analysis indicated diverged in the early 17th century) have an average pairwise percent identity of 77.6% (nucleotide) and 90.1% (amino acid) [10]. These are well within the range for detection by IDseq. Meanwhile, Zika virus, the viral species which caused the recent 2015 epidemic, was discovered in 1940 and shares, on average, 55.6% amino acid sequence identity with dengue virus and 57.0% with West Nile virus [11]. Had the sequence of the first Zika isolate not been in the database, IDseq would not have been able to flag the presence of this virus and researchers would be required to evaluate assembled, unmapped, contigs offline [12].

#### Internal Benchmark Datasets Results

Two additional datasets were simulated to evaluate IDseq’s performance for detection of common clinical microbes (CCM) as well as for disambiguating closely related bacterial species (CRB) (Methods, **Table S3**). The CCM dataset contains reads simulated from 13 total species, including six viral, four bacterial, and three eukaryotic pathogens. The CRB dataset contains reads simulated from 10 species of bacteria, all from the Enterobacteriaceae family. Given that IDseq’s pipeline returns a species-level assignment for each mapped read and the web interface presents results at the species level, we evaluate each of these benchmarks only at the species level. The Kraken2 algorithm assigns reads with ambiguous mappings to higher taxonomic levels. Therefore, the results shown include only the species-specific alignment results.

For the CCM dataset, IDseq filtered out 15.3% of reads during the host and QC filtering pipeline steps. Of the remaining 99,320 non-host reads, IDseq NT showed the greatest per-species recall across the 13 species included (1.0, IQR = 0.99 – 1.0). IDseq NT per-species recall was significantly higher than that of both IDseq NR (0.87, IQR = 0.85 – 1.0, p = 0.02) and Kraken2 (0.89, IQR = 0.64 – 1.0, p = 0.005 Wilcoxon rank sum) (**Figure S3A**). All tools successfully identified the presence of all 13 microbial species. In addition to these species, IDseq NT, NR, and Kraken2 identified 63, 303, and 230 false positive species, respectively. Since the IDseq pipeline returns a species-level assignment for all mapped reads, even in cases where the species may align equally to two different species, it had a notably greater portion of the total (post-qc) reads mapping across those false positive organisms (3.0 % by nt, 10.0 % by NR) than Kraken2, which had only 0.56 % of reads mapping to the false positive species. Kraken2 avoids larger percentages of reads being associated with false-positive species calls by calling a significant portion of ambiguously mapped reads at higher levels of the taxonomic tree.

Similar trends were observed for the CRB dataset. IDseq filtered out 15.6% of reads during the host and QC filtering pipeline steps, leaving 84,410 non-host reads for down-stream analysis. Notably, IDseq nt demonstrates the highest per-species recall (0.68, IQR = 0.43 – 0.80), significantly different than NR (0.46, IQR = 0.15 – 0.61, p = 0.005, Wilcoxon rank sum) and Kraken2 (0.18, IQR = 0.11 – 0.35, p = 0.005, Wilcoxon rank sum) (**Figure S3B**). All tools successfully identified the presence of all 13 microbial species. In addition to these species, IDseq NT, NR, and Kraken2 identified 83, 371, and 167 false positive species, respectively. Again, IDseq NT and NR had greater proportions of total reads mapping to these false-positive species (31.7% and 49.7% for NT and NR, respectively) as compared to Kraken2, with only 0.6 % of reads mapping to false-positive species and the majority of ambiguous reads mapping at higher levels of classification (70.9%).

The impact of ambiguous reads is exaggerated in cases where we simulate reads from multiple closely related species with known genomic similarity. In many cases of an infection, with a single dominant organism, the importance of recall may outweigh the identification of lower-level false-positive species. Additionally, these simulations sample from across the genome. However, we know that ribosomal sequences can be used for typing of bacteria and studies have previously shown improved sensitivity with RNA-seq, where ribosomal RNA comprises the greatest portion of sequenced nucleic acid [13,14]. Thus, IDseq results must be interpreted by the researcher with respect to the sample type and sequencing prep.

## Supplemental Figure

**Figure S1:**
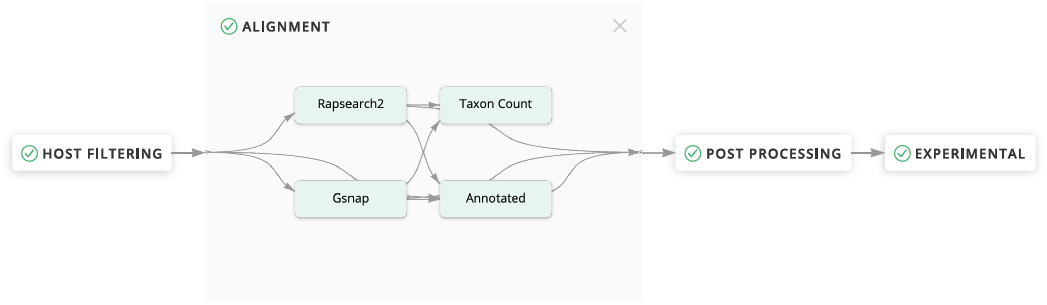
The IDseq pipeline visualization indicates each step in the underlying pipeline and includes a description of the raw command parameters as well as the ability to download intermediate files for offline analysis.

**Figure S2:**
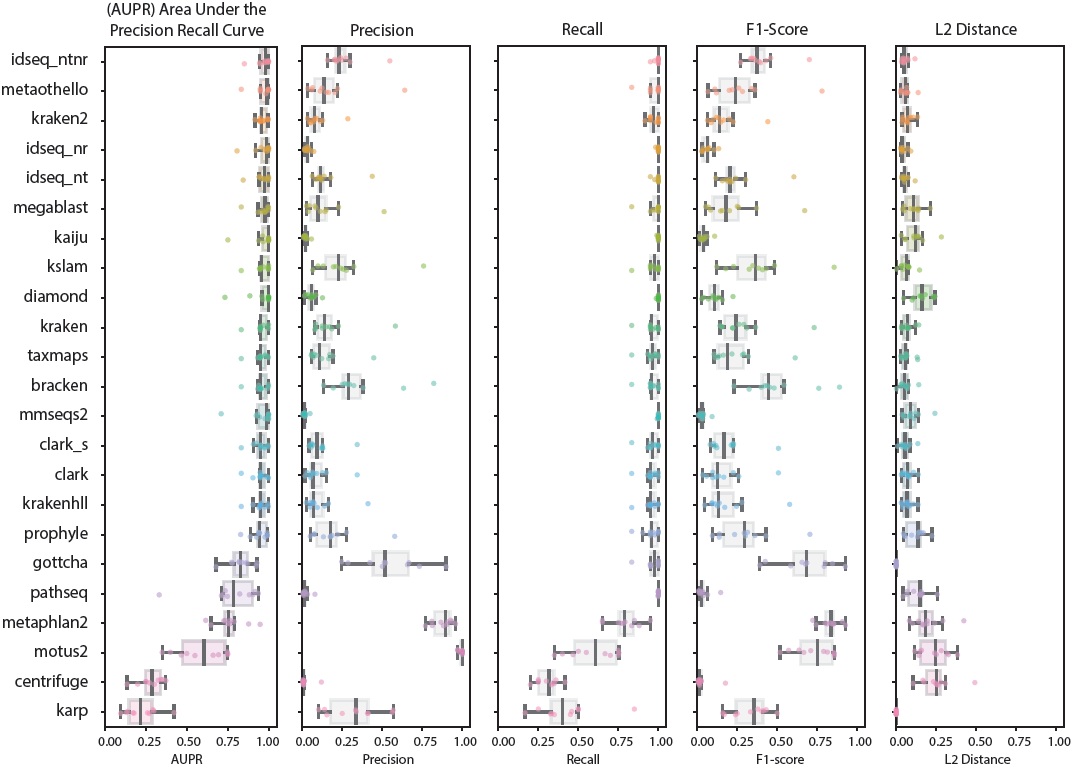
Performance metrics evaluated across 20 mNGS taxonomic identification tools, **A)** Area under the precision recall curve (AUPR), ranges from 0 to 1 (best), **B)** Precision, ranges from 0 to 1 (best). **C)** Recall, ranges from 0 to 1 (best) **D)** F1-score, the harmonic mean of precision and recall values. **E)** L2 distance, ranges from 0 (best) to 1.

**Figure S3:**
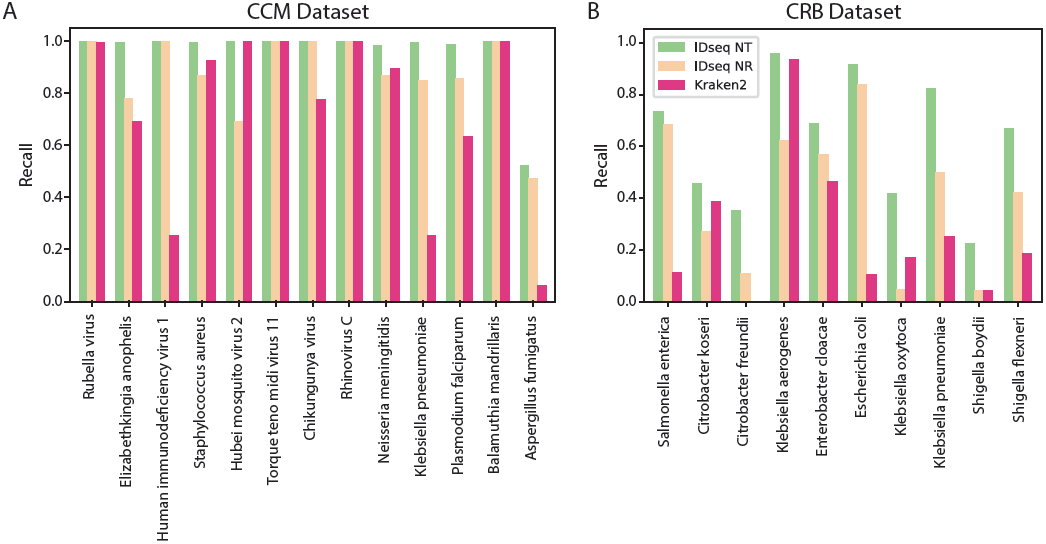
Per-species recall values for two internal benchmark datasets, **A)** The common clinical microbes (CCM) dataset and **B)** the closely related bacteria (CRB) dataset.

## Supplemental Tables

**Table S1:** Github repositories containing open-source code for IDseq pipeline, web application, and benchmarking resources.

| Tool | Code Location |
| --- | --- |
| IDseq Pipeline | https://github.com/chanzuckerberg/idseq-dag |
| IDseq Web Application | https://github.com/chanzuckerberg/idseq-web |
| IDseq Benchmarking Tool | https://github.com/chanzuckerberg/idseq-bench |

**Table S2:**
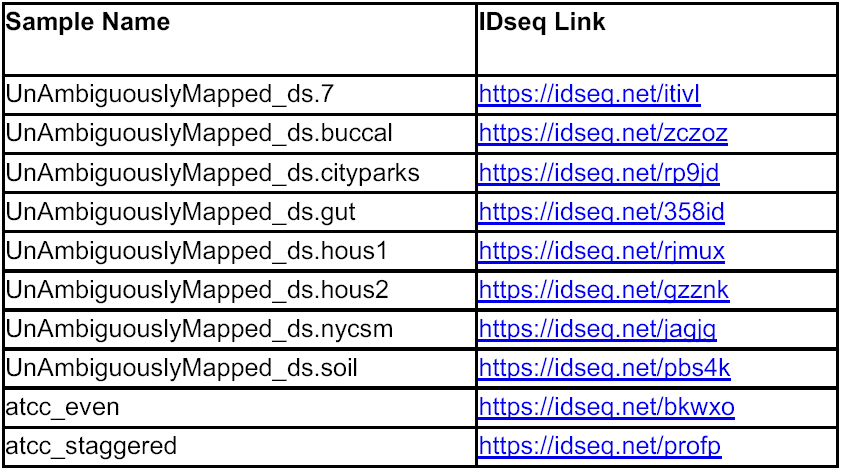
External benchmark datasets and their corresponding IDseq links.

**Table S3:** Internal benchmark datasets and their corresponding IDseq links.

| Sample Name | Project Name | Benchmark Description | IDseq Link |
| --- | --- | --- | --- |
| HRC_100 | HRhinoC Simulation | Divergent Rhinovirus C, 100% identity to reference | https://idseq.net/lx7cf |
| HRC_099 | HRhinoC Simulation | Divergent Rhinovirus C, 99% identity to reference | https://idseq.net/1a3we |
| HRC_095 | HRhinoC Simulation | Divergent Rhinovirus C, 95% identity to reference | https://idseq.net/7a0he |
| HRC_090 | HRhinoC Simulation | Divergent Rhinovirus C, 90% identity to reference | https://idseq.net/8gmj7 |
| HRC_085 | HRhinoC Simulation | Divergent Rhinovirus C, 85% identity to reference | https://idseq.net/q4wks |
| HRC_080 | HRhinoC Simulation | Divergent Rhinovirus C, 80% identity to reference | https://idseq.net/vf8hm |
| HRC_075 | HRhinoC Simulation | Divergent Rhinovirus C, 75% identity to reference | https://idseq.net/qje7h |
| HRC_070 | HRhinoC Simulation | Divergent Rhinovirus C, 70% identity to reference | https://idseq.net/s299t |
| HRC_065 | HRhinoC Simulation | Divergent Rhinovirus C, 65% identity to reference | https://idseq.net/n6ndl |
| HRC_060 | HRhinoC Simulation | Divergent Rhinovirus C, 60% identity to reference | https://idseq.net/46kfh |
| HRC_055 | HRhinoC Simulation | Divergent Rhinovirus C, 55% identity to reference | https://idseq.net/hggw6 |
| HRC_050 | HRhinoC Simulation | Divergent Rhinovirus C, 50% identity to reference | https://idseq.net/a8z8a |
| HRC_045 | HRhinoC Simulation | Divergent Rhinovirus C, 45% identity to reference | https://idseq.net/64ita |
| HRC_040 | HRhinoC Simulation | Divergent Rhinovirus C, 40% identity to reference | https://idseq.net/guat0 |
| HRC_035 | HRhinoC Simulation | Divergent Rhinovirus C, 35% identity to reference | https://idseq.net/shygl |
| HRC_030 | HRhinoC Simulation | Divergent Rhinovirus C, 30% identity to reference | https://idseq.net/hdc6r |
| HRC_025 | HRhinoC Simulation | Divergent Rhinovirus C, 25% identity to reference | https://idseq.net/v84fc |
| CCM Dataset | Benchmark v1 | Common clinical microbes, simulated with idseq-bench | https://idseq.net/ak8qu |
| CRB Dataset | Benchmark v2 | Closely related bacterial species, simulated with idseq-bench | https://idseq.net/71wns |

